# Unraveling Endothelial Cell Phenotypic Regulation By Spatial Hemodynamic Flows With Microfluidics

**DOI:** 10.1101/2020.08.28.268599

**Authors:** Sarvesh Varma, Guillermo Garcia-Cardena, Joel Voldman

## Abstract

Human endothelial cells (hECs) experience complex spatiotemporal hemodynamic flows and that directly regulate hEC function and susceptibility to cardiovascular disease. Recent medical imaging studies reveal that helical flows strongly correlate with lowered disease susceptibility, as contrasted to multidirectional disturbed flows. However, a lack of platforms to replicate these spatial profiles of flow (SPF) has prevented biological studies to investigate the role hECs play in tuning the observed SPF-correlated disease susceptibility. Here, we utilize microfluidic devices to apply varying SPF upon hECs for the first time, and discover that these flows can differentially impact hEC morphology, transcription, and polarization. Collectively, our platform and studies significantly advance our ability to delineate flow-regulated hEC function and disease susceptibility.

**Significance Statement:** *In vivo*, hECs experience complex hemodynamic flows, including those that are spatially helical or disturbed, which is in stark contrast to the unidirectional flows typically used to study hECs *in vitro.* Understanding the impact of SPF on hEC function informs our understanding of the pathophysiology of hEC dysfunction and can lead to interventional solutions that specifically perturb SPF to lower disease risk. Here, we leverage microfluidics to apply and discover the specific impact of SPF on hECs for the first time. Broadly, our platform bridges the mutual interests of the vascular biology and interventional cardiology communities to collectively understand how cardiovascular health is tied to the way blood flows upon the endothelium.

## Introduction

Blood ejected from the heart traverses upon the vascular endothelium as complex motion: where its spatial trajectories emerge from the interplay of blood rheology and vessel architecture; and where its temporal velocities emerge from pulsatile contractions generated with each heartbeat. Notably, human vascular function as well as risk of cardiovascular disease are regulated by the way blood traverses on the endothelium (1). Specifically, arterial flow exhibiting a high fluid shear stress (FSS), pulsatile waveform, and with unidirectional (1) or helical (2) spatial profile of flow (SPF) is associated with functional endothelium *in vivo*. However, perturbation of this SPF to one with a low-and-oscillatory FSS waveform and a multidirectional (so-called disturbed) trajectory (as occurs near vascular branches) causes endothelial dysfunction (3, 4). Although it is well established that endothelial function is broadly coupled to flow *dynamics*, the impact of *SPF* on endothelial cell phenotype is still largely unknown (5).

Understanding how endothelial cell function is affected by SPF is of importance to both interventional cardiologists and vascular biologists. In these pursuits, each community utilizes tools (e.g. stents, perfusion systems) that perturb flow profiles experienced by endothelial cells. Strutted stents or grafts are known to induce local disturbed flow that increases the risk of restenosis, thrombosis and neointimal hyperplasia (6), whereas flow helicity has been postulated to reduce flow disturbances and impede atheroprone flow signatures (7-9). Indeed, specific stents have been used to emulate physiological helical flows (7-9), and a recent clinical trial has demonstrated improved vessel patency, and therefore the clinical utility of reproducing helical SPF (10).

The loss of flow helicity has also been correlated to the development of human cardiovascular diseases, such as atherogenesis, pulmonary hypertension, and atheromatous renal vascular diseases (11-13) and thus flow helicity has been postulated as an atheroprotective feature in several arterial regions (2, 14). To support this argument, current hypotheses in the vascular computational fluid dynamics (CFD) community revolve around the effect of helicity on mass-transport of cells, lipids, oxygen, etc.(2, 15).

Despite the clinical and CFD evidence of the disease-protective nature of helical SPF and disease-initiating nature of disturbed SPF, there have been no experimental models to specifically study such SPF in the context of vascular biology. Here we leverage a microfluidic platform able to create uniform, helical, and chaotic flow to characterize—for the first time—the specific coupling of SPF to human endothelial cell (hEC) phenotype. We designed grooved microchannels to generate uniform, helical and chaotic SPF with identical average FSS and temporal waveforms, and then used the platform to examine the influence of SPF on hEC morphology (alignment and elongation), junction integrity, transcriptional regulation (KLF2 expression; NRF2 and NF-κB activity) and planar cell polarity. Our results demonstrate that, even in the absence of the temporal waveforms, SPF can *differentially* regulate hEC biology at the single cell- and population level. These results should now motivate the community to consider SPF as a first-order factor (along FSS magnitude and temporal profile) regulating hEC biology.

## Results

### Microfluidic device design and flow validation

To emulate helical and disturbed flow found *in vivo*, we patterned microchannels with grooved surfaces(16) on the channel ceilings (Fig. 1A). In the microfluidics community, grooved microchannels have primarily been designed and optimized for improved fluidic ‘mixing’ within laminar flow, but have not, to our knowledge, been used to generate SPF for cell cultures. We adapted the grooved microchannels by volumetric scaling of the original SG (staggered groove) and SH (staggered herringbone) devices (16) for adequate media volumes and for patterning ECM coatings (17),(18). With these changes, we could routinely obtain similar healthy pre-flow monolayers in our devices to a static dish-cultured controls.

**Figure 1.**
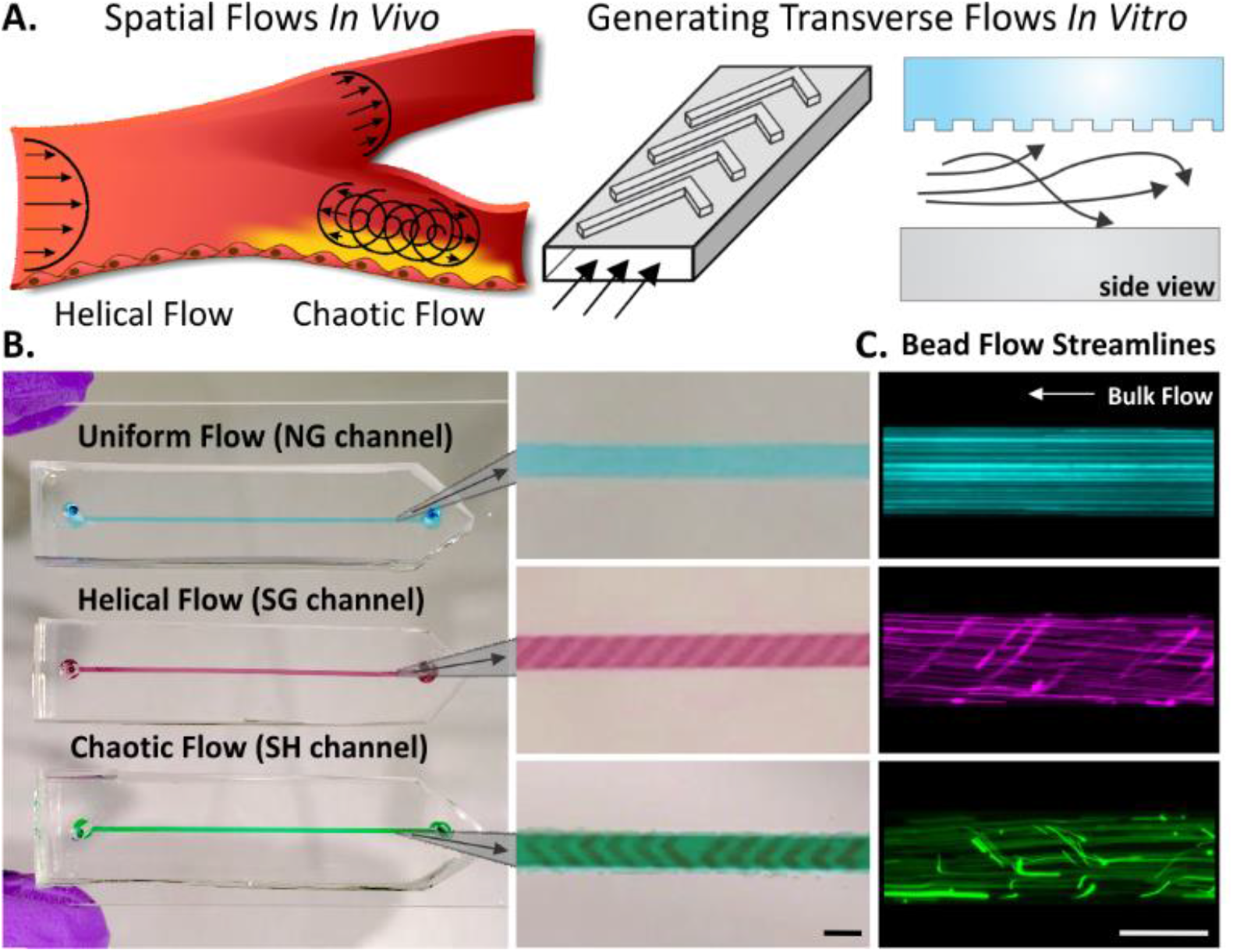
Emulating *in vivo* spatial profiles of flow within microfluidic devices. **A.** Helical and chaotic SPF are found upon atheroprotected and atheroprone vascular regions *in vivo*. **B.** Grooved-ceiling microfluidic channels generate predefined 3D transverse flows within in the laminar flow regime. **C.** Image of the NG SG and SH devices with colored fluids and close-up of the groove architectures in the respective channels. Fluorescent 4.5 µm bead flow streamlines generated at 20 dynes/cm^2^ validate uniform flow in NG channels, helical flow in SG channels and chaotic flow in SH channels. Scale: 0.5 mm.

To generate helicity, we designed SG that can introduce left- or right-handed helical flows. The groove depth ratio to the channel height, aspect ratio, groove angle and orientation were all designed (19) to achieve predictable localized normalized helicity(20) (LNH), which is a standard metric for measuring helicity in the vascular CFD community. LNH is a spatially-varying but time-averaged metric for helicity (SI Appendix, Fig. S1). For unidirectional flow, LNH = 0, while LNH=1 is maximal (right-handed) helicity. We used CFD modeling (similar to our previous work (21)) to design SG with LNH up to ∼0.7 (average ∼ 0.3), which is desirably in the range of the reported *in vivo* helicity (LNH: 0.2-0.8) (20).

To mimic the disturbed behavior of *in vivo* SPF, we incorporated SH grooves (16) into the microchannels. SH grooves induce steady chaotic flow (22, 23), with an average LNH ∼ 0, indicative of non-helical multidirectional flow, as desired. Notably, the chaotic flow in SH channels refers to the fluid particle pathlines in such channels, and will result in a wall FSS that varies over space but not time (for steady applied pressure). We realized that due to their inherent micromotion and migration (ubiquitous for *in vitro* cultures), hECs would effectively sample the spatially-varying SPF in SH channels, converting it to a time-varying SPF from the cells’ perspective. Although, spatial sampling also exists in helical flow (SG) channels and in uniform channels without grooves (no groove channels, NG), the FSS profiles experienced in those channels are steady and spatially invariant. This feature allows us to study the relative impact of SPF upon hECs within these channels.

We fabricated channels of each type (Fig. 1B) and observed fluorescent bead streamlines consistent with each flow type (Fig. 1C) and with the well-understood fluid mechanics expected in each channel.

### Structural Adaptations to the Endothelium Induced by Helical and Chaotic SPF

We first wanted to determine whether hECs could even sense and respond differentially to different SPF, which would be required for regulating endothelial function and ultimately disease susceptibility. With this scope, we chose to first investigate how the *presence* of helicity (as opposed to the *degree* of helicity) affects hECs. We exposed hECs to constant physiological FSS of 20 dynes/cm^2^ (temporal average of characteristic atheroprotective waveform (24)) to decouple the impact of flow dynamics on hECs (which has been widely studied, in the absence of SPF) from that by the SPF (which largely remains unknown). We used human umbilical vein endothelial cells (HUVECs) as these hECs have been widely used for decades by the vascular biology community to study hEC mechanotransduction. Importantly, despite their venous origin, studies have shown that differences between the arterial and venous are dependent on the *in situ* conditions (e.g. flow-induced arterialization of venous grafts), instead of being fixed by cell fate decisions (25).

Under our device perfusions, hECs could be cultured for at least 1 week, where they maintained expected flow-induced alignment (e.g. parallel alignment in NG channels), as well as continuous junctions (marked by VE-Cadherin staining, Fig. 2A). In contrast, similar cultures exposed to helical SPF (SG channels) demonstrated homogenous helical alignment (Fig. 2B). To the best of our knowledge, these results are the first demonstration of applying helical flows on hECs *in vitro* and observing helical alignment similar to what has previously only been observed *in vivo*(26, 27). Alignment was unaffected by cell seeding density within a range of 30,000 cells/cm^2^ (i.e. 40% of our target seeding density, SI Appendix, Fig. S2), suggesting that the alignment was specific to the applied SPF. Conversely, cells exposed to chaotic SPF generally did not align (Fig. 2B). Additionally, helical and uniform flows induced actin stress fibers aligned to the flow direction, whereas cells under chaotic SPF generally induced fewer axial, and more cortical, distributed stress fibers, similar to static conditions (Fig. 2B).

**Figure 2.**
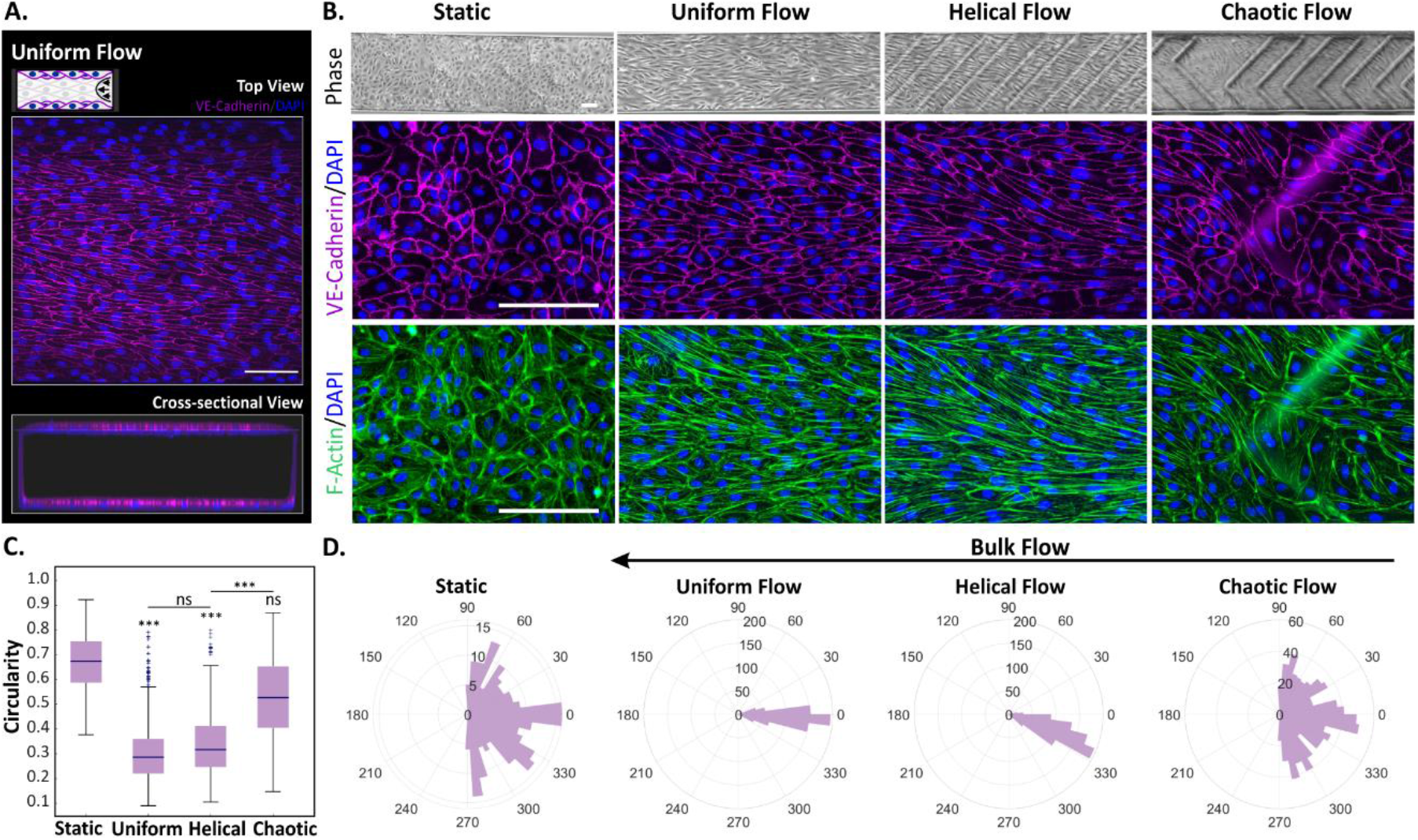
Assessment of endothelial morphological adaptations to SPF. **A.** Illustration of healthy hEC culture under physiological FSS (20 dynes/cm^2^) in NG channels after 1 week of perfusion. Top- and cross-sectional confocal images exemplify continuous VE-Cadherin junctions and nuclear staining within the channels. **B.** Cellular junctions, elongation, alignment are shown via phase microscopy and immunofluorescence imaging of VE-Cadherin and cytoskeletal stress fibers (F-Actin staining). These morphological adaptations are differentially induced in hECs exposed to uniform, helical or chaotic SPF (20 dynes/cm^2^, 72h). **C.** Cellular aspect ratio is quantified by circularity in the same flow conditions against static controls (n = 3, 725 cells/condition; ***: p<0.001). **D.** Distribution of cellular alignment in uniform, helical and chaotic SPF against controls, (n = 3, 725 cells/condition). Scale: 0.1 mm.

We further quantified cellular circularity (Fig. 2C) and observed that, while in static conditions hECs maintained a cobblestone-like morphology (high circularity, ∼0.67), both helical and uniform flows statistically decreased circularity, as cells elongated in the flow direction. While helical and uniform SPF elongated cells in a statistically indistinguishable manner, chaotic SPF relatively inhibited hEC elongation (Fig. 2C). Cells under chaotic SPF had a higher degree of circularity compared to the helical flows (p < 0.001). A study by Levesque *et al.* showed that if aortic stenosis was introduced surgically upon an otherwise atheroprotected endothelial region *in vivo*, then the stenosis-induced multidirectional flows caused elongated and aligned ECs to increase their circularity and lose their orientation to the overall flow direction(28). Although it is challenging to attribute this morphological change specifically to the flow directionality *in vivo*, our experimental results show that similar morphological changes can be induced in hECs exposed to multidirectional flows in our devices.

When we quantified alignment distributions in the different SPF (Fig. 2D), we found that uniform and helical SPF induced similar alignment histograms, though cells in helical SPF were aligned with the direction of the helicity rather than the channel direction. In contrast, despite a mean alignment towards bulk flow, cells in chaotic SPF were more similarly distributed to static controls (SI Appendix, Table SI). These results demonstrate—for the first time—that hECs exhibit distinct morphologies under helical and chaotic SPF in the absence of pathogenic stimuli, and thus that hECs can respond to different SPF.

### Impact of Helical and Chaotic SPF upon Transcriptional Regulators of hEC Phenotype

To understand how the morphological changes connect to the underlying transcriptional program, we examined regulation of Kruppel-like factor 2 (KLF2), which is widely regarded as a critical regulator of hEC function to sustain atheroprotection(3, 4). While several studies have investigated the role of hemodynamic waveforms on KLF2 expression, the impact of SPF on its regulation has not been explored. Hence, we asked whether KLF2 expression was modulated by these distinct flow SPF by using a KLF2-GFP transcriptional reporter (Fig. 3A)(29).

**Figure 3.**
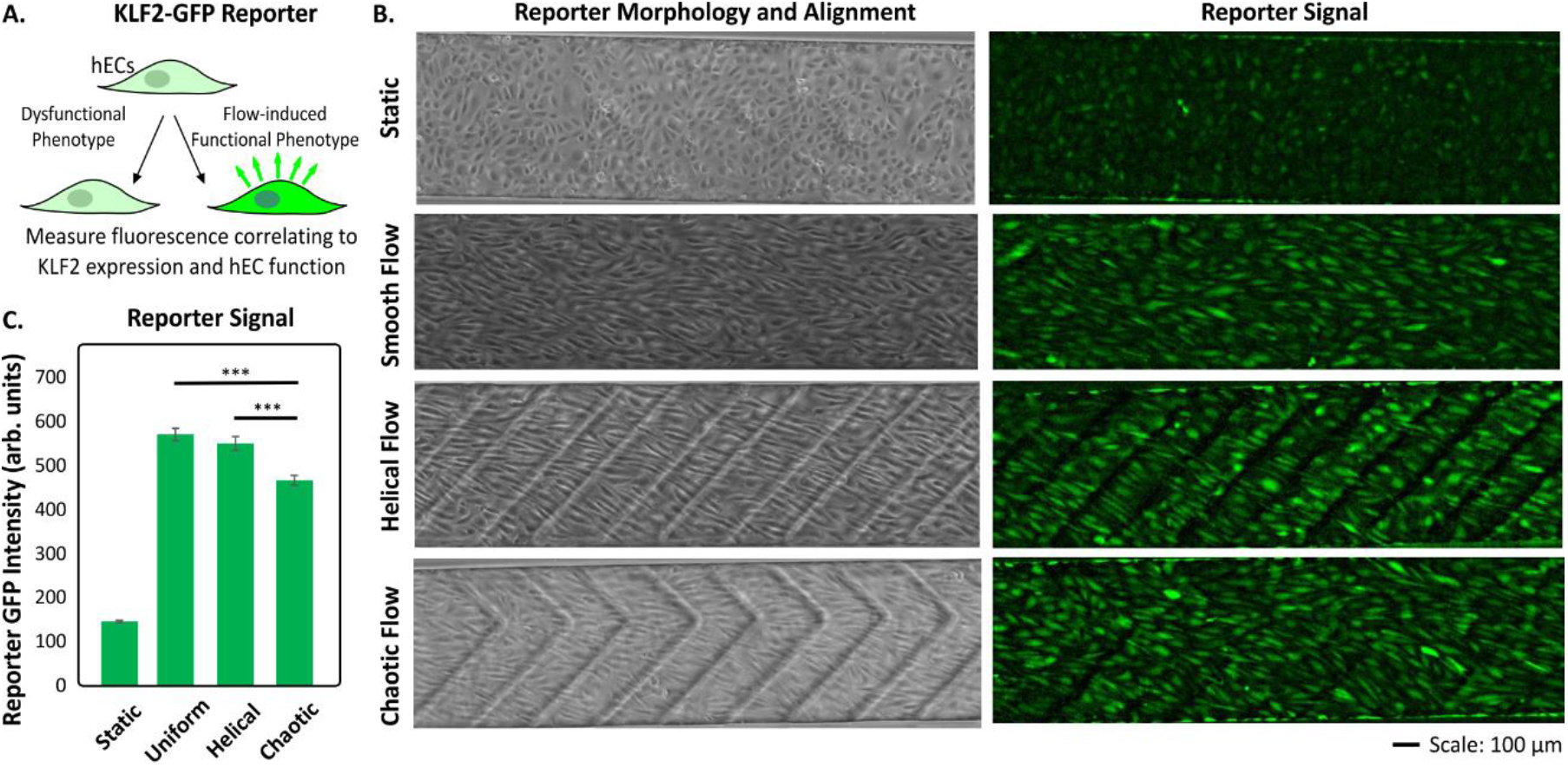
Assessment of KLF2-GFP reporter response to SPF. **A.** Endothelial KLF2-GFP FSS reporter activity is tied to KLF2 expression, and thus serves as a surrogate readout for the induction of a functional phenotype. **B.** Reporter morphology and reporter GFP signal observed under uniform, helical or chaotic flow with constant FSS of 20 dynes/cm^2^, 48h against static controls. **C.** Reporter responses (population median GFP) measured by quantitative imaging and shown for each flow condition against static controls (n ∼1000 cells/condition, 6 independent devices, same culture passage, error: +/-SEM, ***: p<0.001).

We applied helical, chaotic and uniform SPF at 20 dynes/cm^2^ for 48h to hEC reporter cultures. We observed enhanced reporter induction to all flow stimuli compared to static controls (Fig. 3B). Reporter induction under chaotic SPF was not completely surprising, since the high applied FSS magnitude could activate the reporters regardless of the SPF. However, we found a statistically significant (p<0.001) decrease in the reporter response when cells were exposed to chaotic SPF (3.2 fold) as opposed to helical SPF (3.8 fold), thus revealing an inhibitory effect of chaotic SPF on transcriptional regulation (Fig. 3C). We further validated these expression trends using an independently generated batch of KLF2-GFP reporters (SI Appendix, Fig. S3). KLF2 expression has been shown to upregulate FSS-induced actin fiber assembly though RhoA, while suppression of KLF2 inhibits hEC alignment to flow(30), providing a putative mechanism to connect morphology associated differences to KLF2 expression under the two SPF.

We also investigated whether SPF impacted other transcriptional regulators (i.e. NF-κB and NRF2), by assessing their sub-cellular localization as a readout for their activity (SI Appendix, Fig. S4). ECs exposed to atheroprone dynamics acquire a pro-inflammatory phenotype characterized by NF-κB pathway activation, adhesion molecules expression, and KLF2 suppression (24). In contrast, flow-regulated nuclear translocation of NRF2 evokes an anti-oxidative phenotype *in vitro* and *in vivo*(31, 32). Previous *in vitro* studies show p65 translocation to the nucleus within ∼min upon constant, uniform SPF(33); within ∼min upon oscillatory, uniform SPF(34), and upto ∼24h upon atheroprone, uniform SPF(24). In contrast, *in vivo* studies report p65 remains ‘primed’ in the cytosol of ECs even in atheroprone regions of murine and porcine models, in the absence of risk factors *in vivo*(35, 36*). Hence, whether multidirectional SPF per se* could cause sustained p65 or NRF2 translocation remained unknown. Here, our results show that distinct SPF induce NRF2 translocation but do not cause sustained p65 translocation, which corroborates with ‘steady state’ observations seen *in vivo*. Collectively, as activation of NF-κB pathway and the NRF2 antioxidant pathway, were not differentially affected, SPF was selective in its effects on regulatory networks.

### Helical and Chaotic SPF Differentially Regulate Endothelial Planar Polarity

We next examined EC planar cell polarity (PCP), which is a hallmark of the functional hEC phenotype that critically regulates physiological functions and has previously been associated with regions of varying SPF. For example, EC PCP is lost in the atheroprone region of the aortic arch and is gradually established in neighboring ECs experiencing atheroprotective flow (37), suggesting a link between the local spatiotemporal flow to the PCP. *In vivo* imaging, meanwhile, has shown that ECs polarize and helically migrate upstream of the blood flow during development (38), and surgically induced stenosis caused ECs in the multidirectional flow region to reverse their polarity (39). We used the microfluidic platform to determine whether SPF *per se* could induce changes in PCP.

We first imaged hECs under uniform SPF (SI Appendix, Fig. S5). We quantified the angle of the golgi complex (GM130) relative to the nucleus, as well as to the bulk flow, to classify single cell polarization (37, 40) as a readout for PCP (SI Appendix, Fig. S6). Within the first 24h, a larger fraction of hECs had their golgi and microtubule organizing center (MTOC, γ-Tubulin) polarized downstream and migrated downstream, consistent with prior seminal studies (39, 41, 42). After sustained perfusion (between 46-68h), however, a subpopulation of hECs started migrating upstream, and at 72h nearly half of hECs (49.1 +/- 3% SEM) polarized upstream. Such upstream polarization has also been observed in murine capillary venous cells and zebrafish intersegmental venous cells, as well as human coronary arterial ECs *vitro* (43). As this emergent behavior reflected the upstream polarization levels measured *in vivo* (e.g., during vessel development *(44)*), we chose to assess hEC PCP 72h after initiation of perfusion.

Without flow, golgi and nuclei orientations were widely distributed (Fig. 4A-C). However, with uniform and helical SPF, the cell bodies and nuclei aligned parallel and helically, respectively, with a narrower distribution of angles. Helical flow also induced predominant upstream polarization (53.5 +/- 5% SEM) as compared to uniform flow. In contrast, chaotic SPF induced wide distributions in nuclear and golgi alignment, effectively resulting in a reduced polarization. In addition, we found that these polarization patterns persisted in chronic application (1 week) of SPF and that chaotic SPF again impeded polarization (SI Appendix, Fig. S7). These results demonstrated a stark difference between hECs conditioned with helical and chaotic SPF, and that overall, helical and chaotic SPF can markedly impact hEC PCP profiles.

**Figure 4.**
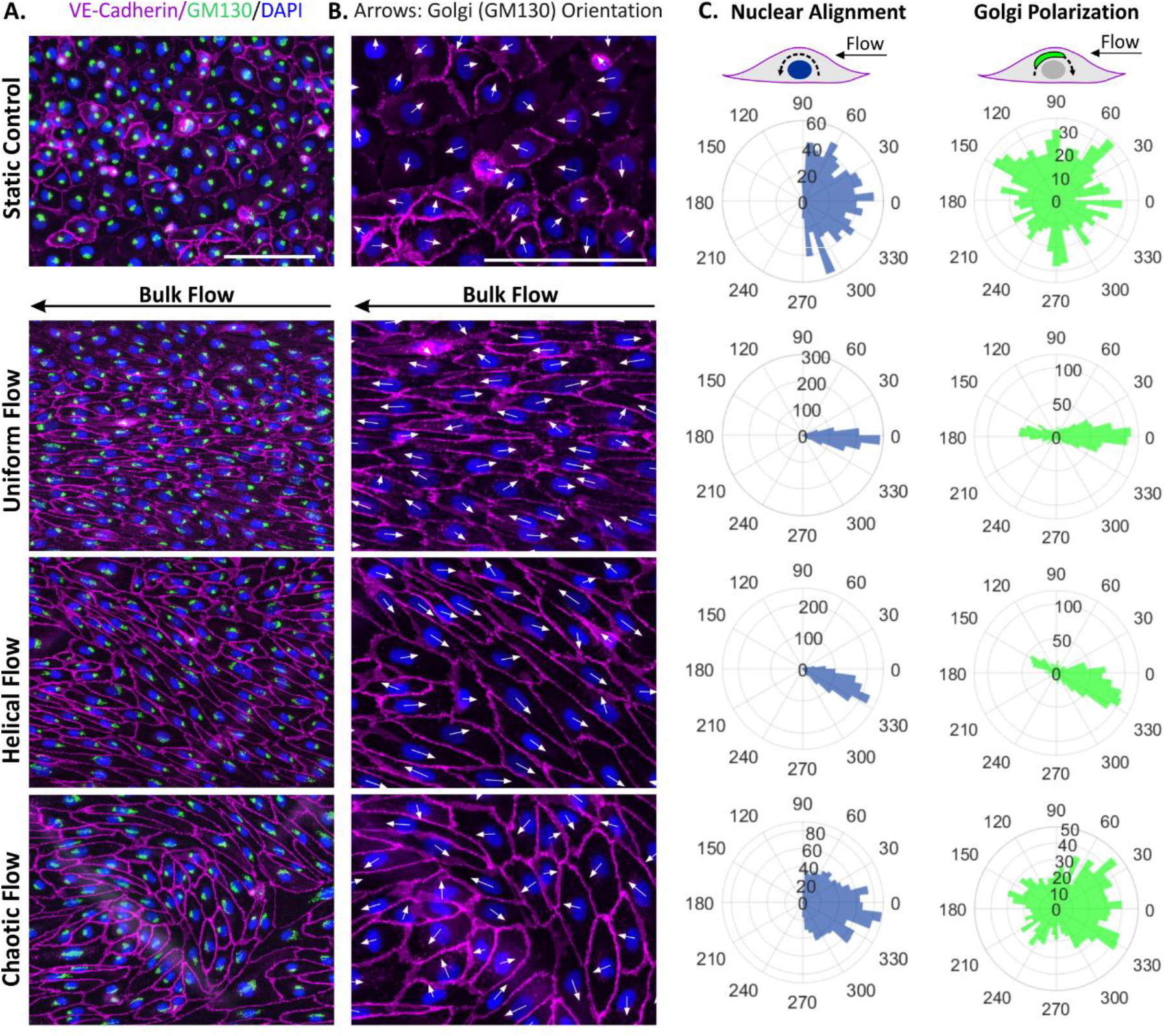
Establishment of endothelial planar cell polarity under distinct SPF. **A.** Subcellular localization of hEC golgi complex (GM130) in hECs exposed to static (no flow), uniform, helical, chaotic SPF at (20 dynes/cm^2^, 72h). **B.** Arrows indicating golgi orientation in each hEC. Images are shown from one representative experiment. **C.** Nuclear alignment and golgi polarization distributions are shown under the same conditions. (Data pooled from 3 independent experiments, 1250 cells/condition). Scale: 0.1 mm.

As the ‘random’ golgi polarization under chaotic SPF resembled the static condition at the population level, we asked whether individual hECs polarize to chaotic SPF in the same way as if there was no flow. Examining the nuclear and golgi orientation coupling on a per-cell level, in static culture for any given nuclear angle the golgi tended to be oriented ∼90° to the nuclear axis (Fig. 5A). hECs conditioned to uniform or helical SPF had nuclei orientations clustered around the expected cell orientation angle, similar to Fig. 4, and the golgi angle clustered either around +/-180° (downstream) or around 0° (upstream), without apparent coupling around 90° (unpolarized state). In contrast, hECs exposed to chaotic SPF lost the golgi angle coupling of ∼90° observed under static controls, and instead retained a statistically different median polarization angle and overall angular distribution (p< 0.001, SI Appendix, Table SII). Thus, chaotic SPF specifically decoupled the nuclear-golgi orientation in comparison to other SPF, illustrating that these hECs were likely phenotypically distinct.

**Figure 5.**
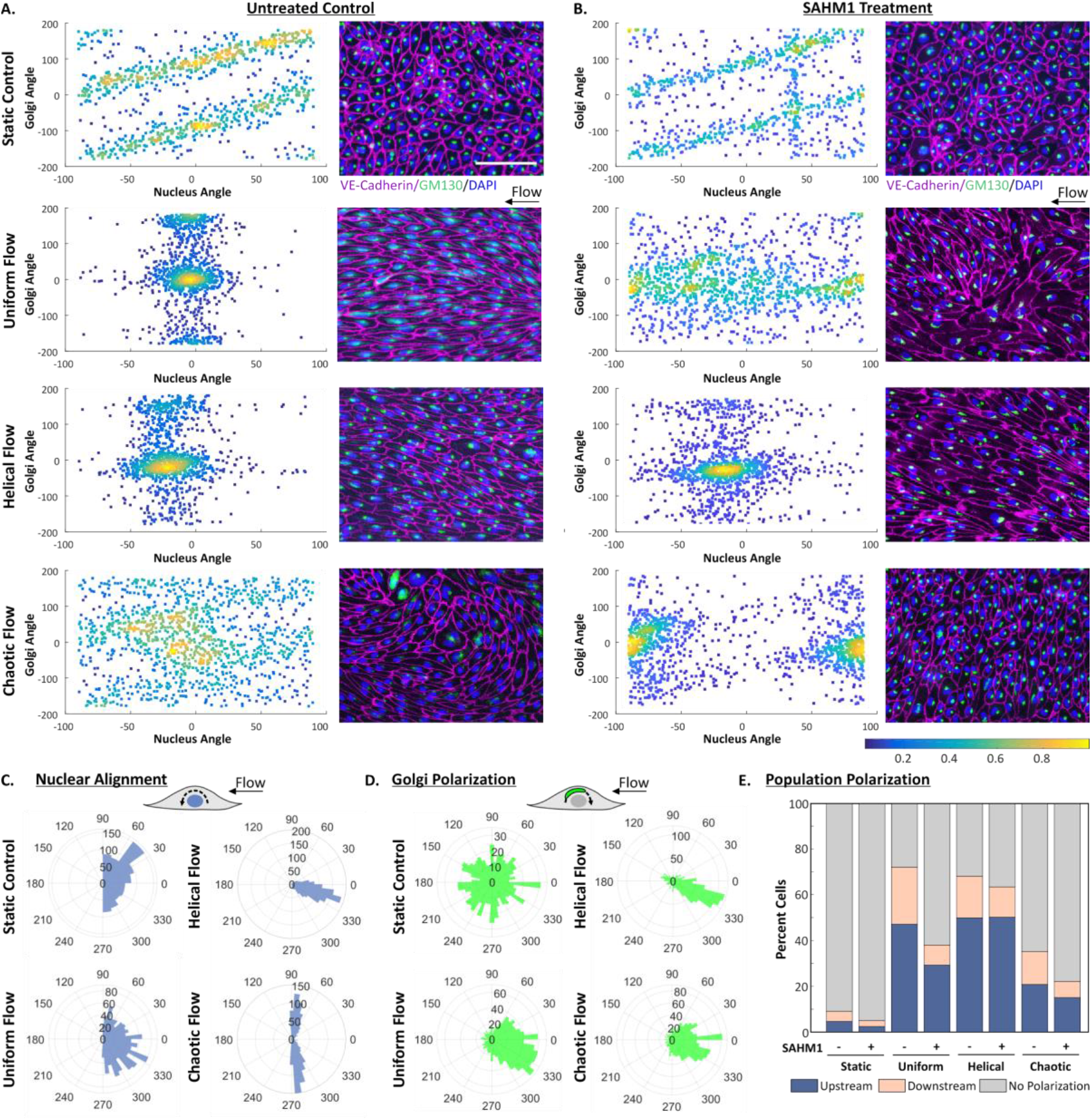
Coupling of nuclear and golgi orientation and overall polarization at the single cell level under distinct SPF. Density scatter plots of the nuclear and golgi orientation for single cells under distinct SPF, without (**A.**) or with (**B.**) SAHM1 treatment. Subcellular localization of hEC golgi complex (GM130, green stain) with respect to the cell (VE-Cadherin, magenta stain) and nucleus (DAPI, blue stain) are shown as well. All data was accumulated at 72h after perfusion at 20 dynes/cm^2^. Control cells were preconditioned with flow for 48h, followed by vehicle treatment and the NOTCH1-inhibited cells were similarly preconditioned and treated with SAHM1 at 10 µM for 24h under flow. **C.** Nuclear alignment and (**D.**) golgi polarization distributions are shown under the same SAHM1-treatment conditions. **E.** Population polarization was determined based on single-cell polarity classification, with or without SAHM1 treatment (for all panels: n =2, 1250 cells/condition). Scale: 0.1 mm.

Since polarization and morphology both responded to distinct SPF, we hypothesized that helical and chaotic SPF may be differentially sensed by the EC mechanosensors. We focused on NOTCH1, which is an essential regulator of EC junctional integrity, alignment and elongation, as well as of PCP and migration (45-47). Inhibition of NOTCH1 signaling has been shown to adversely alter hEC polarization to flow (45). Although recent studies highlight the necessity of NOTCH1 in establishing *de novo* PCP (45) the role of NOTCH1 in PCP established under helical and chaotic SPF remains unexplored. Hence, we tested the putative role of NOTCH1 in the maintenance of PCP, by preconditioning hECs with distinct SPF (48h) and subsequently inhibiting NOTCH1 within the same flow conditions.

We blocked NOTCH1 by direct inhibition of its transcription complex (48) through SAHM1 treatment (Fig.5), and in separate experiments by inhibition of upstream γ-secretase signaling (45, 47) through DAPT (SI Appendix, Fig. S9). Under static conditions neither treatment adversely affected morphology (SI Appendix, Fig. S8) or nuclear and golgi polarization (Fig. 5B, SI Appendix, Fig. S9), however both treatments repressed NOTCH1 transcription and pathway associated genes (SI Appendix, Fig. S8), as expected. In our flow experiments, both SAHM1 and DAPT treatment inhibited nuclear and golgi angle coupling and increased the fraction of unpolarized cells in static controls and in all SPF conditions (Fig. 5D, E, SI Appendix, Fig. S9), although to varying degrees depending on the SPF preconditioning. Specifically, SAHM1 perfusions revealed unique polarization responses and nuclear-golgi coupling in hECs preconditioned with distinct SPF (Fig. 5B, SI Appendix, Fig. S10). For instance, NOTCH1 inhibition disrupted the angle coupling patterns observed in hECs preconditioned with uniform SPF (Fig. 5A, SI Appendix, Fig. S9A) and broadly resulted in the loss of cellular, nuclear and golgi alignment to flow (Fig. 5C,D, SI Appendix, Fig. S9C,D). In contrast to uniform or chaotic SPF, hECs preconditioned with helical SPF resisted depolarization by SAHM1, as reflected by the population median and distribution analyses (Fig. 5B, SI Appendix, Fig. S10, Table SIII). Furthermore, analysis of coupling distributions (SI Appendix, Fig. S11) revealed that inhibiting NOTCH1 through DAPT in uniform SPF preconditioned hECs caused the golgi-to-nuclear polarity coupling to phenocopy that in hECs under chaotic SPF in a statistically significant manner, while the same treatment under helical SPF failed to do the same with significance (SI Appendix, Fig. S11, Table SIII). Although hEC responses to DAPT and SAHM1 were not identical (possibly due to differences in molecule targets and potency), our correlative results show that broadly, hECs could polarize differentially as a consequence of their prior conditioning by distinct SPF.

Interestingly, NOTCH1 inhibition also increased the cellular (and nuclear) tendency to align perpendicular to the bulk flow direction in all treatment conditions (Fig. 5C, E, SI Appendix, Fig. S9), which is consistent with observations by Mack *et al.*, where NOTCH1 inhibition caused ECs to align perpendicular to flow *in vitro* and *in vivo* (45). Overall, our results on hEC PCP resonate with our conclusions on hEC morphological adaptations and transcriptional regulation, in that SPF is a distinct biomechanical cue for hEC phenotype, alongside the well-appreciated flow dynamics and intensity (FSS).

## Discussion

Due to the established causal nature of hemodynamics in human cardiovascular pathophysiology, studying the spatiotemporal complexity of blood flows is of long-standing interest to communities as diverse as vascular biology, computational fluidics dynamics, and interventional cardiology. Quantitative imaging of human hemodynamics has revealed that its complexity emerges from preexisting combinations of flow parameters upon disease-protected and disease-prone vascular regions, including FSS magnitude, dynamics and directionality. Out of these multiparametric combinations, we know with certainty that that low-and-oscillatory magnitude FSS primes ECs towards a dysfunctional state, while high-pulsatile magnitude FSS enhances EC function towards a functional state (1). Moreover, there is now also an accumulation of observational evidence linking flow directionality (specifically helical and multidirectional chaotic SPF) to cardiovascular disease susceptibility (2, 49). However, to-date existing *in vitro* models have been incapable of reproducing such SPF, and hence there have been no biological studies that elucidate whether ECs can sense and respond to these SPF *per se*.

Mimicking the *in vivo* SPF *in vitro* has been an ongoing challenge for both macroscale and microscale flow platforms. For instance, to mimic disturbed flows, ECs have been exposed to turbulence within macroscale platforms (50) *in vitro*. Although turbulent flow exhibits chaotic characteristics, turbulence itself does not occur in physiological vessels *in vivo* (51). At hemodynamic Reynolds numbers, SPF within disease-prone vessel regions (52) is more accurately described as spatially unsteady and chaotic (53, 54), which is indeed reproducible in our devices. Realizing FSS gradients are indeed part of the *in vivo* flow complexity, prior platforms have generated spatial gradients of FSS and have provided to unique insight of EC responses to spatial cues of flow (55-57). However these models still do not recapitulate the helical or chaotic directional profiles observed *in vivo*. Furthermore, in these approaches each cell receives a distinct spatial shear stimulus, making it is difficult to decouple emergent behaviors at the single cell level to that of the collective heterogeneous population. Motivated by the lack of technological solutions to mimic *in vivo* SPF, we leveraged microfluidics to investigate the phenotypic regulation of hECs by helical and chaotic SPF for the first time.

ECs *in vivo* experience the entire spatiotemporal complexity of hemodynamic blood flow. This makes it impossible to exclusively link the *spatial* aspect of the blood flow to the phenotypic response. In this regard, our approach presents a unique advantage in that it allowed us to selectively apply helical and chaotic SPF (found in the two regions), and importantly, under identical, time-invariant FSS of physiological magnitude. Our results broadly demonstrate that in addition to differentially impacting hEC morphology and alignment, helical and chaotic SPF differentially polarize hECs in a NOTCH1-dependent manner. Notably, we discovered that at the subcellular level, the coupling of golgi alignment to nuclear orientation differs according to SPF and NOTCH1 inhibition.

One of the first descriptions of SPF was made in the 1500s by Leonardo da Vinci in his drawings of spiraling flows from the aorta (5, 58). In the 1960s, flow vortices were first reproduced and explained using models of aortic valves (59). Subsequent studies reported helical alignment of endothelial cells *in vivo* (27), helical arrangements in the sub-endothelial cells and matrix (60), and helical migration of ECs during development (38). Here, microfluidic devices designed to create different flow patterns allowed us to demonstrate that helical and chaotic SPF can drive hEC phenotype. While our results do not definitively demonstrate that helical or chaotic SPF alter disease susceptibility, our experiments now link previously reported observations from animal models to human ECs. Further, beyond the use of microfluidics to compare distinct types of SPF, these types of microfluidics devices can be extended to systematically explore quantitative features of SPF (such as the degree of flow helicity) through design variations, as well as enable future studies that reveal the coupling of temporal FSS dynamics and SPF to hEC biology and the underlying molecular mechanisms. These tools now bridge the mutual research interests between the vascular biology community and computational fluid dynamics community to collectively understand the how human cardiovascular health is tied to the way blood traverses upon the vascular endothelium.

## Materials and Methods

### Microfluidic device fabrication

The microfluidic culture devices were cast in PDMS from custom-made plastic replica molds using standard soft-lithography protocols. The devices constituted of a cell culture chamber of dimensions: 35 mm length x 0.5 mm width x 0.2 mm height, with 0.1 mm grooves, designed according to our previous work(21). Each device was plasma-bonded to a borosilicate glass slide and all surfaces were chemically cross-linked with a 0.1% gelatin solution, per our previously published protocol(17). Briefly, following surface conjugation of glutaraldehyde, a 0.1% aqueous gelatin solution was introduced for 30 min and subsequently, cell culture media was introduced into each chamber to prime the device. The entire disconnected device was stored in a cell culture incubator for 16-24h to equilibrate the device with media constituents and the cell culture environment.

### Computational Fluid Dynamics Simulations and Flow Validation

Flow simulations were carried out in the COMSOL Multiphysics (Burlington, MA) environment with the particle tracing module. The simulations were carried out similar to those described previously(21). Briefly, channels geometries were imported and the channel flow velocity, and derived LNH values were simulated using with a constant flow rate determined according to the desirable channel FSS. Channels set to be 500 µm wide, 250 µm high, with a groove-to-channel height ratio (termed α) of 0.4 for LNH quantification. Particle tracing was performed using 2 μm particles in SG and SH channels were calculated with a net forward, constant inflow applied at a wall FSS of 20 dynes/cm^2^. Wall FSS (and associated flow conditions) in our devices was calculated analytically with conditions and assumptions outlined in our previous work(18), and was validated with numerical simulations conducted in COMSOL. At 20 dynes/cm^2^, the channel Reynolds number for our experiments was ∼37. Although we used the same molds for soft-lithography based fabrication, there can still be small structural variances in the channel created during assembly which, in addition to any unintentional temporal instabilities of flow, could collectively affect the flow profiles in the same device over experimental repeats. As it is challenging to experimentally measure such variances at our Reynolds number flows in 3D, particle flow streamlines were qualitatively assessed by recording fluorescent bead flow in the NG, SG, and SH channels. For channel flow validation, 4.5 µm fluorescent beads were suspended in PBS buffer at a concentration of 500,000/mL, and were perfused through each device at a wall FSS of 20 dynes/cm^2^. Given the significant differences in the channel architectures, we expectedly never observed unintentional flow disturbances in one device that would resemble flow profiles expected in the other channels. Importantly, our studies focused on the relative differences between uniform, helical and chaotic SPF which were reproducibly generated in the respective channels, as per our simulations and qualitative observations and matched the profiles well established in the literature.

### Cell culture and treatments

Primary human umbilical endothelial cells (HUVEC) were isolated and cultured in the Garcia-Cardena Lab (Brigham and Women’s Hospital, Boston, MA) according to standard procedures. These cells were cultured and maintained in EGM-2 Bulletkit (CC-3162, Lonza). KLF2-GFP reporter cells were cultured M199 medium (12117F, Lonza) containing 20% fetal calf serum, 1% L-glutamine, 1% penicillin/ streptomycin, 100 μg/ml heparin and 50 μg/ml endothelial cell growth supplement (BT-203, Biomedical Technologies). The KLF2-GFP reporters are defined as two batches of KLF2-GFP cells generated from distinct pooled HUVEC populations, and characterized according to the published methods(29). Device perfusions were carried out using equilibrated medium for the respective cell populations described above. All cells were cultured on tissue culture polystyrene dishes treated with 0.1% v/v gelatin solution (ES-006-B, Millipore) and were incubated in a humidified incubator at 37°C and 5% CO2 until they reached confluence. All experimentation was carried out within the first 10 passages of the culture. For Notch inhibition experiments, perfusion medium was supplemented with 10 µM DAPT (S2215, Sellekchem) or 10 µM SAHM1 (491002, Sigma) after 48h of flow conditioning, with controls supplemented with vehicle (DMSO). Such a treatment method was chosen to allow cells to first adapt to the flow onset, and to the SPF.

### Gene expression Analysis

Cellular mRNA was extracted (RNeasy Micro kit, Qiagen74004), quantified, reverse-transcribed (ProtoScript II First Strand cDNA Synthesis Kit, NEB E6560) and used for quantitative real-time PCR (with iQ SYBR Green Supermix, Bio-Rad 170-8882) on the Bio-Rad CFX96 system, all according to the recommended protocols. Gene expression profiles were quantified and normalized relative to GAPDH expression and primer sequences are provided in Supplemental Table IV.

### On-chip Perfusion

After priming devices with medium, hECs were seeded into devices at a seeding density of 75,000 cells/cm^2^. Confluent cultures and continuous junctions were routinely observed within 4-6h, after which the culture monolayer was washed with fresh medium and connected to the flow setup. A closed-loop peristaltic pump (RP-1, Rainin Instrument) was setup with a flow dampener and used in accordance to our previous work(18). Flow rates were set in accordance to the calibrated values and were verified prior to running experiments. Auxiliary fluidic connections and tubing were autoclaved, assembled, and equilibrated with culture medium overnight in the incubator along with devices, 24h prior to cell seeding. Similarly, the entire medium perfusion volume was equilibrated prior to experimentation and kept in the incubator with tubing routed to devices for perfusions.

### Real time Imaging and cell tracking

For on-stage perfusions and live-cell imaging, devices were kept under a custom built isolated microenvironmental enclosure that enabled equilibration of humidified 5% CO2. In these experiments, medium was supplemented with 25 mM HEPES. Images were acquired every 30 minutes using customized MATLAB scripts under the phase channel (for assessing hEC morphology) of a Axiovert 200M microscope (Zeiss) fitted with a cooled CCD camera ImagerQE (LaVision) and an automated stage Ludl MAC 5000 (Ludl) using 5-20x objectives. Cell tracking was performed using the MTrackJ plugin in ImageJ and each track analysis was conducted in MATLAB.

### Immunofluorescence Reagents and Staining

On-chip staining was conducted by adapting the manufacturer’s protocols for respective reagents. Cells were fixed with 10% formalin (HT5011, Sigma Aldrich) for 10 minutes, washed twice with PBS, permeabilized with 0.1% Triton-X 100 solution for 10 minutes (X100, Sigma Aldrich) in 1× PBS and washed twice again with PBS. Cells were then blocked with 1% BSA (15260037, Life Technologies) diluted in PBST (28352, Life Technologies) for 2 hours and were washed with PBS. Primary antibodies (1:200 dilution) were incubated at 37°C in blocking buffer for 3h and secondary antibodies applied for 1 h at room temperature. Primary antibodies used in the work include: goat anti-human CHD5 (AF938, Novus Biologicals), mouse anti-human γ-Tubulin (sc-17788, SCBT), mouse anti-human p65 (sc-8008, SCBT), rabbit anti-human NRF2 (sc-722, SCBT), rabbit anti-human GM130 (EP892Y, Abcam). Secondary antibodies include: Alexa Fluor 555 donkey anti-mouse IgG (A31570, Life Technologies), Alexa Fluor 488 donkey anti-rabbit IgG (A21206, Life Technologies), Alexa Fluor 647 Donkey Anti-Goat IgG (705-605-147, Jackson ImmunoResearch), all used at 1:400 dilution. In some cases, cells were also counterstained with NucBlue (R37606, Life Technologies) and ActinGreen 488 Ready Probes (R37110, Life Technologies), according to the manufacturer’s protocols.

### Quantification of Planar Cell Polarity and Reporter Signals

For KLF2-GFP reporter signal quantification, cells were imaged with Zeiss Axiovert 200m with a mercury lamp excitation light source, with 5x objectives using a standard GFP filter (ex:470/40, em:525/50 nm). Cells were also stained and imaged with a live-cell nuclear marker (Nucblue, Life Technologies) or whole-cell marker (Celltracker, Life Technologies) that clearly identified each cell in separate fluorescence channels at the end the experiment, according to the manufacturer’s protocols. The cell masks were transferred to the background-corrected FITC images to quantify single cell responses. For every device, at least 500 and typically 1000 cells were quantified and their median GFP fluorescence was either plotted as-is, or normalized to the signals from static controls. Images were analyzed in ImageJ (NIH), and region of interests were defined by automated masking were verified prior to quantification. Image analyses and data visualization plots were carried out using custom MATLAB scripts. To quantify cell polarity, immunofluorescence images of GM130, DAPI and VE-Cadherin channels were background-corrected and imported to ImageJ, where DAPI and VE-Cadherin stains were used to determine nuclear and cellular ROI and properties using custom ImageJ macros. VE-Cadherin stains were used to determine cell morphological features, whereas nuclear masks were used to determine nuclear morphological features (e.g. alignment to flow). Following individual cellular and nuclear annotation, each cell was associated with its GM130 stain and the object centroid was connected to that of the nucleus to define Golgi directional vector for each cell. Dividing cells, or those partially in the field of view were discarded from analyses. For each experimental condition, 3-5 random fields of views were analyzed, and data from independent experiments was pooled for analyses. Subsequently, paired angular annotations of the nucleus and golgi were exported to MATLAB for data analysis and visualization.

### Statistical analysis

Two-sided Wilcoxon rank sum test was used to compare at independent flow conditions to each other, or against static controls. Statistical analyses were conducted in MATLAB. Angular distributions were compared through their median value by using the non-parametric Kruskal-Wallis H (one-way ANOVA on ranks) test. The distributions themselves were compared by using the Kuiper test. Further details are provided in the Supplementary Information document. Error values are reported as mean ± SEM. For all analyses: *: p-value < 0.05; **: p-value < 0.01 ***: p-value < 0.001.

## Acknowledgments

SV was supported by the MIT-GSK Gertrude B. Elion Research Fellowship. The authors would also like to acknowledge Bendix R Slegtenhorst (Laboratory for Systems Mechanobiology, Center for Excellence in Vascular Biology, Department of Pathology, Brigham and Women’s Hospital, Boston, MA, USA) for generating the KLF2-GFP reporters used in this study. The authors would also like to thank Prof. Roger D. Kamm (MIT) and Prof. Abraham D. Stroock (Cornell University) for their thoughtful feedback on our work.

