## Supplemental Info for "Unraveling Endothelial Cell Phenotypic Regulation By Spatial Hemodynamic Flows With Microfluidics"

#### SI Appendix


##### Supplementary Materials


Fig. S1. Computational fluid dynamics simulation of SPF within SG and SH channels.
Fig. S2. Impact of seeding density upon hEC alignment under distinct SPF.
Fig. S3. Assessment KLF2-GFP sensor responses to SPF.
Fig. S4. NRF2 and p65 activity in response to distinct SPF.
Fig. S5. Monitoring establishment of hEC planar cell polarity under uniform SPF.
Fig. S6. Classification of single cell planar polarity.
Fig. S7. Establishment of hEC population polarity under chronic application of distinct SPF.
Fig. S8. Cellular responses to SAHM1 or DAPT doses under static conditions.
Fig. S9. Coupling of nuclear and golgi orientation under distinct SPF and following DAPT perfusion.
Fig. S10. Cumulative distribution functions and Kuiper tests of relative golgi angles caused by SAHM1.
Fig. S11. Cumulative distribution functions and Kuiper tests of relative golgi angles caused by DAPT.

Table S1. Statistical measurements and tests for cellular orientation.
Table S2. Statistical measurements and tests golgi orientation.
Table S3. Statistical analyses of relative golgi angles under distinct SPF conditions.
Table S4. Primer Sequences for qRT-PCR

### Supplementary Figures

#### Computational Fluid Dynamics

Quantifying Helicity $LNH(\mathbf{x},t) = \frac{\mathbf{v}(\mathbf{x},t)\cdot\boldsymbol{\omega}(\mathbf{x},t)}{|\mathbf{v}(\mathbf{x},t)|\,|\boldsymbol{\omega}(\mathbf{x},t)|}$ $\mathbf{v}(\mathbf{x},t)$ : velocity $\boldsymbol{\omega}(\mathbf{x},t)$ : vorticity LNH> 0: Right-handed flow LNH< 0: Left-handed flow

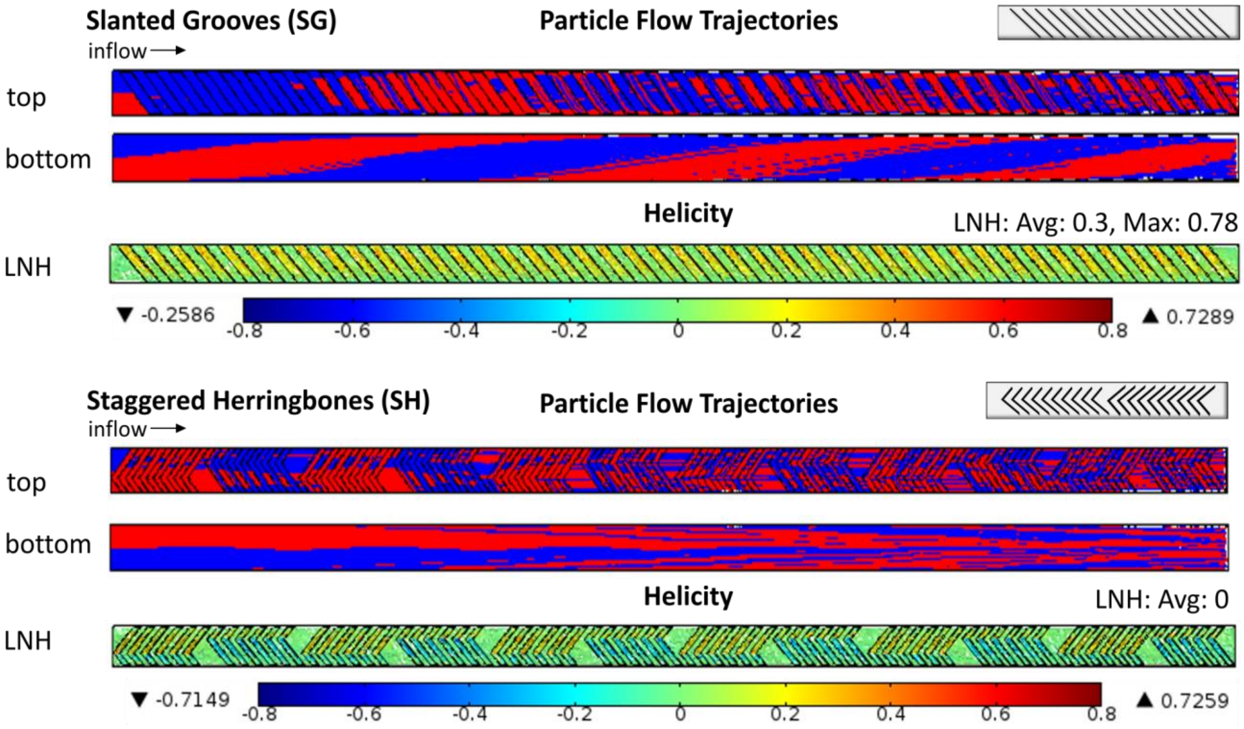


channel width: 500 μm, α: 0.4, channel height: 250 μm, FSS: 20 dynes/cm$^2$ ; 2 μm particles

**Figure S1. Computational fluid dynamics simulation of SPF within SG and SH channels.** CFD modeling of particle flow profiles (red and blue) shows helical flow created in SG channels and chaotic flow in SH flow channels respectively. Flow helicity is quantified by deriving spatial LNH profile in the SG and SH channels and assessing the volumetric average over the entire channel.

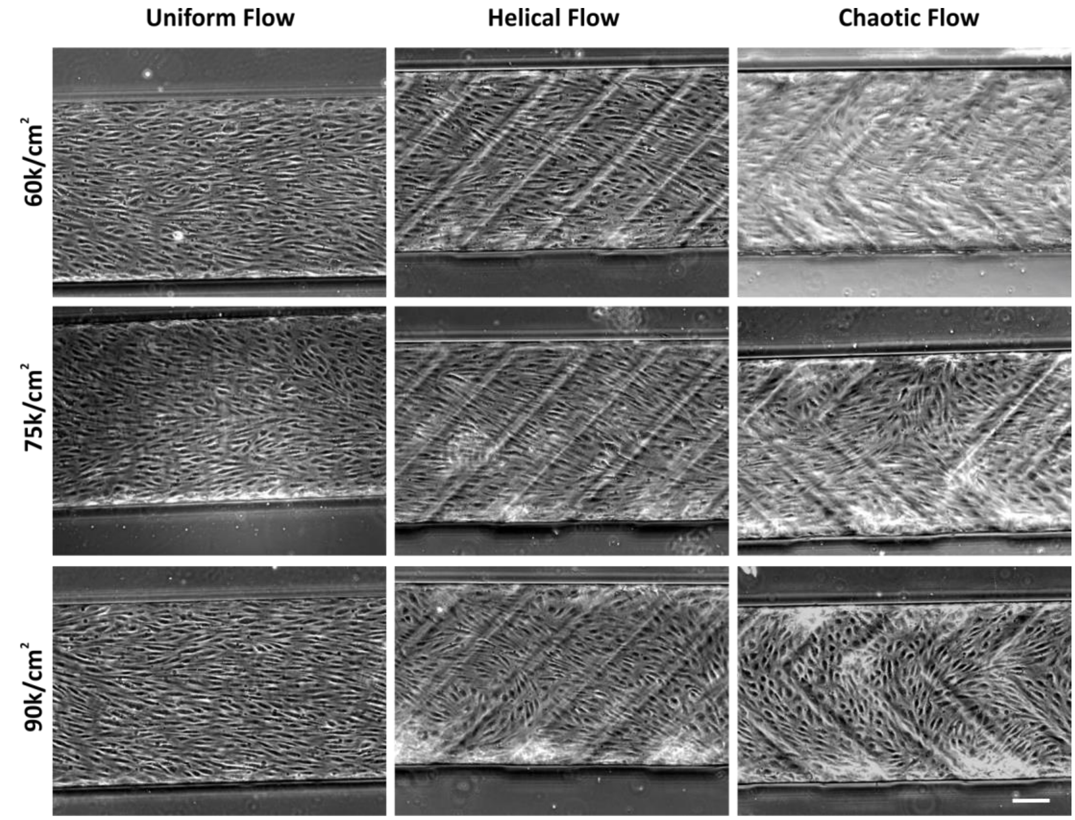


**Figure S2. Impact of seeding density upon hEC alignment under distinct SPF.** hECs were seeded at the shown densities and perfused at a time-invariant FSS of 20 dynes/cm$^2$ for 3 days and fixed prior to imaging. Phase micrograph images of cell alignment under uniform SPF shows parallel alignment with the flow direction; helical alignment in the SG channels and a random alignment under chaotic flows generated in the SH channels, regardless of the seeding densities. Scale bar: 0.1 mm.

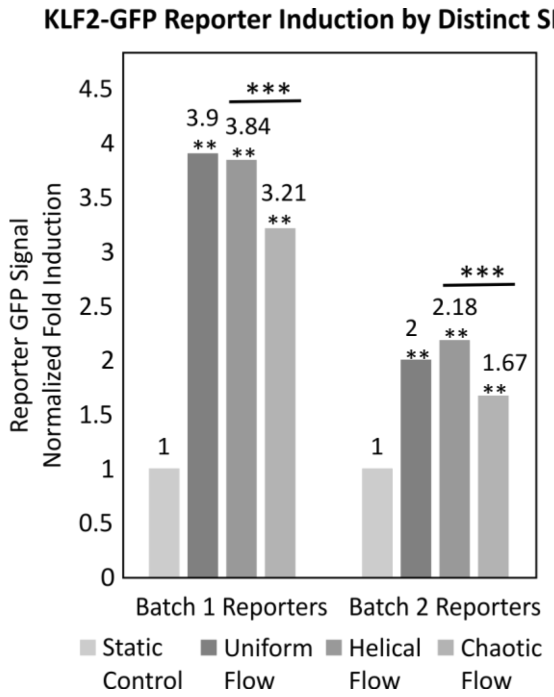


**Figure S3. Assessment KLF2-GFP reporter activity in response to SPF.** Static-normalized KLF2-GFP reporter activity (population median fluorescence) under uniform, helical or chaotic flow with constant FSS (20 dynes/cm$^2$, 48h) profiles of flow against the static control, from two independently generated batches of the reporters (n > 1000 cells/condition, pooled from two independent experiments. ns: not significant; ** p<0.01; *** p<0.001).

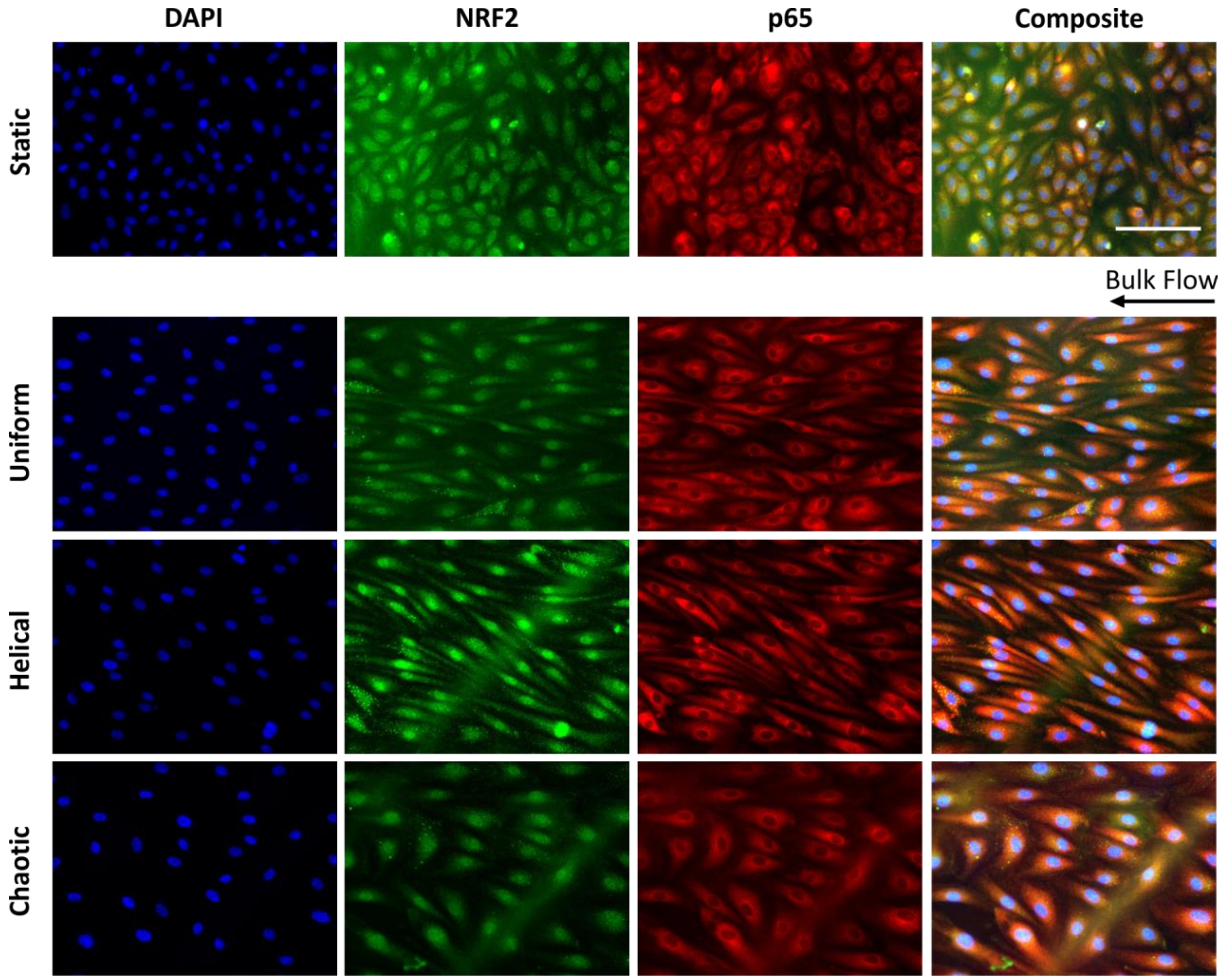


**Figure S4. NRF2 and p65 activity in response to distinct SPF.** hECs were subjected to uniform, helical or chaotic flows at 20 dynes/cm$^2$ for 48h and were stained with antibodies against p65 and NRF2. In contrast to the staining profiles seen in static controls, all flow conditions induced a nuclear translocation of NRF2 (indicative of an antioxidant state). Additionally, similar to static controls, none of the flow conditions caused any translocation of p65 (indicative of an anti-inflammatory state). Scale: 0.1 mm.

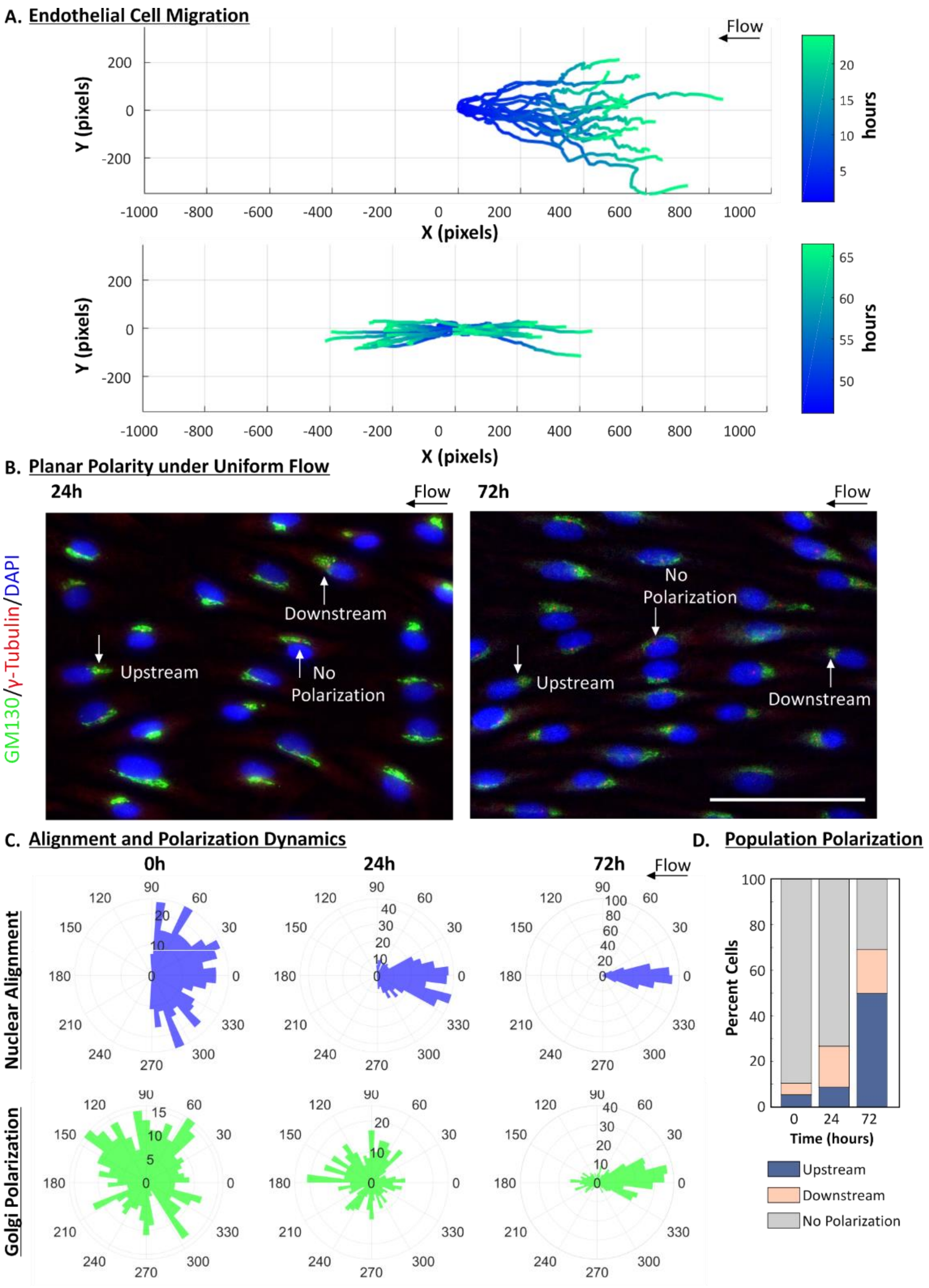


**Figure S5. Monitoring establishment of hEC planar cell polarity under uniform SPF. A.** Migration is quantified for hECs experiencing uniform flow at 20 dynes/cm$^2$ between 0-24h (n= 30 cells), and between 46-68h (n = 60 cells) from a representative experiment. **B.** Planar polarity under the same flow conditions is illustrated at 24h and 72h by staining of GM130 (golgi) and ɣ-Tubulin (MTOC) with a nuclear counterstain. White arrows indicate that the golgi and MTOC can polarize in the same direction within a cell. This direction be in up- or downstream directions to the flow, or be approximately perpendicular to the flow direction (i.e. not polarized) in distinct cells. **C.** Quantification of the planar cell polarity at a population level through the nuclear and golgi alignment before flow, and 24h and 72h after flow (n = 450 cells from 2 independent devices). **D.** Measurement of relative populations from C., that are polarized upstream, downstream, or are not polarized at the same time points. Scale: 50 µm.

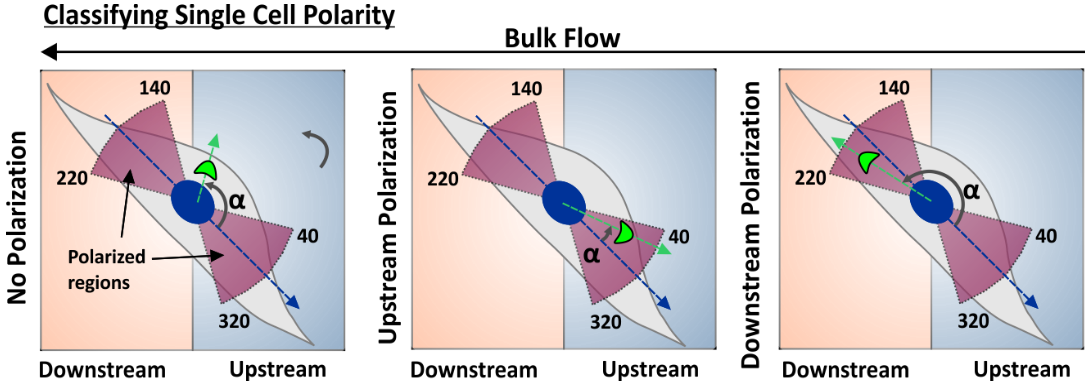


**Figure S6. Classification of single cell planar polarity.** The relative angle of the golgi centroid to the nuclear orientation (α) is measured for each cell and assessed whether it lies within +/- 40 degrees of the nuclear orientation (polarized regions) in the front or back of the nucleus. These locations determine whether the cell is polarized up- or downstream. If the golgi vector lies outside this region, it is classified as not polarized. In all cases, bulk flow is applied from right to left.

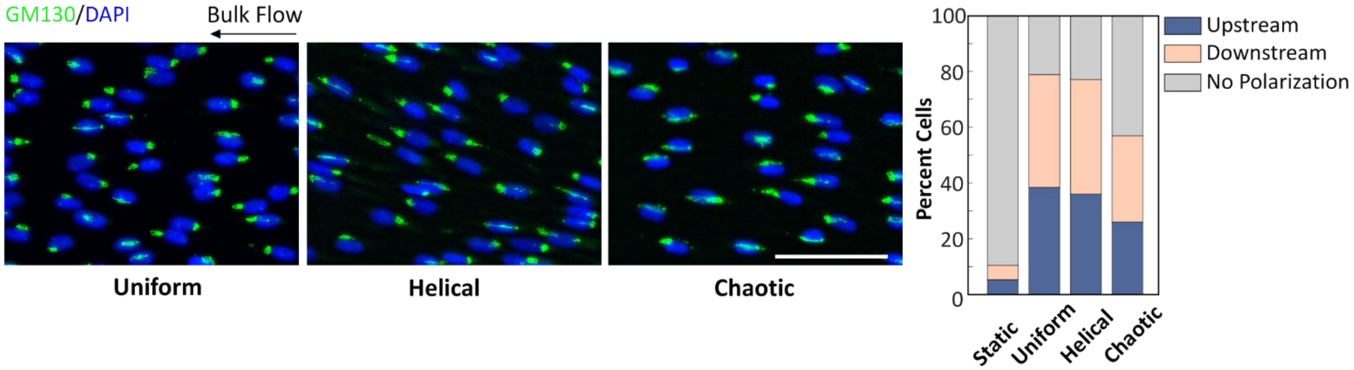


**Figure S7. Establishment of hEC population polarity under chronic application of distinct SPF.** Staining profile of the golgi in hECs exposed to uniform, helical or chaotic flow at 20 dynes/cm$^2$ for 1 week. Cellular polarization was classified as upstream, or downstream, or as not polarized based on the relative angle of the golgi to the bulk flow direction, as described in the methods. Fraction of cells polarized in the three classes under distinct SPF (n = 480 cells/condition pooled from 2 devices). Scale: 50 µm.

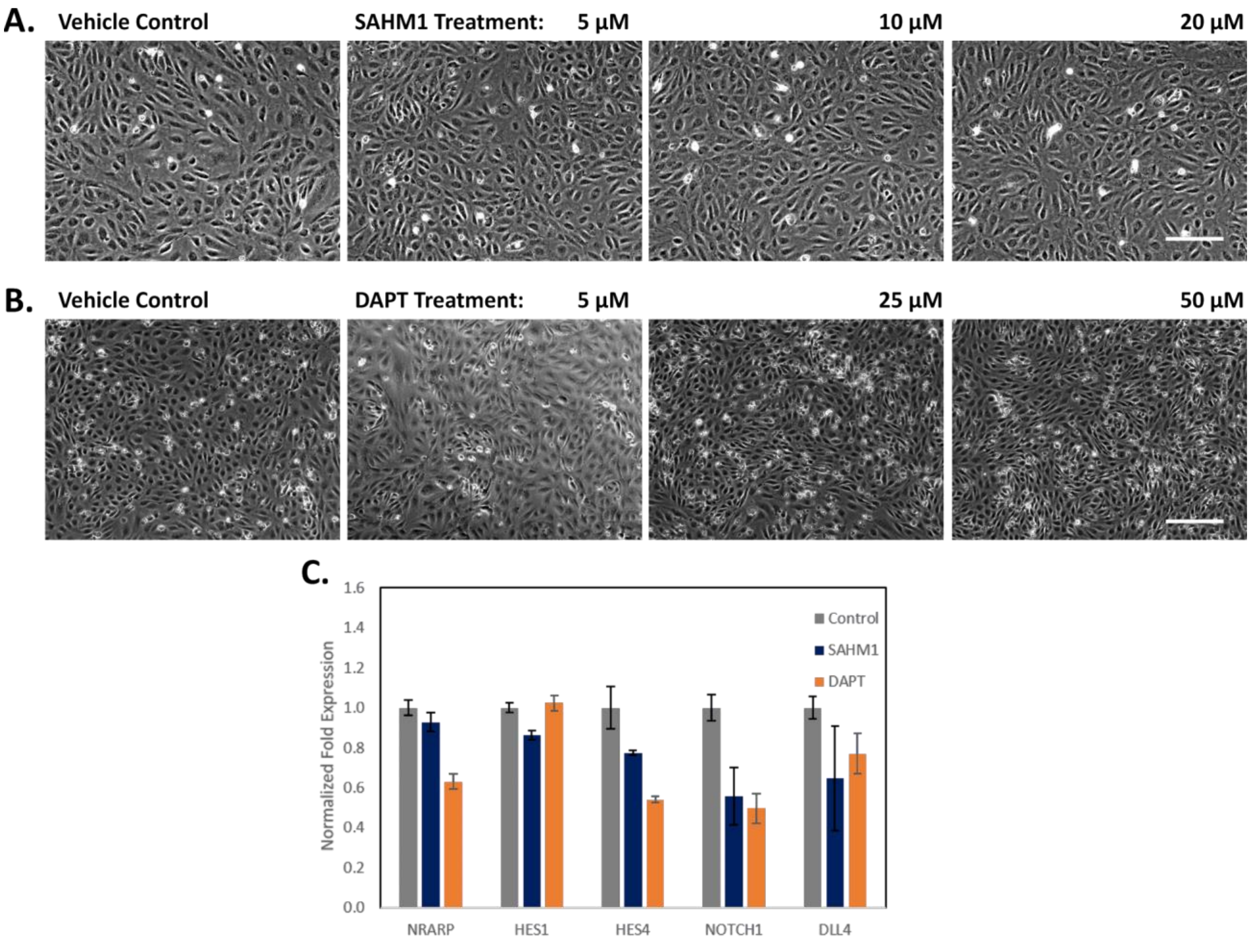


**Figure S8. Cellular responses to SAHM1 or DAPT doses under static conditions. A.** hECs morphology was used as a qualitative assessment of distinct doses of SAHM1 or (**B.**) DAPT, compared to that of the vehicle control (DMSO). Cells were imaged without washing, 24h following treatment. **C.** Normalized fold expression of NOTCH1, and its pathway-associated genes in hECs treated with SAHM1 (10 µM, 24h) or with DAPT (10 µM, 24h). Data averaged from two independent treatments, error: +/- SEM. Scale: 0.1 mm.

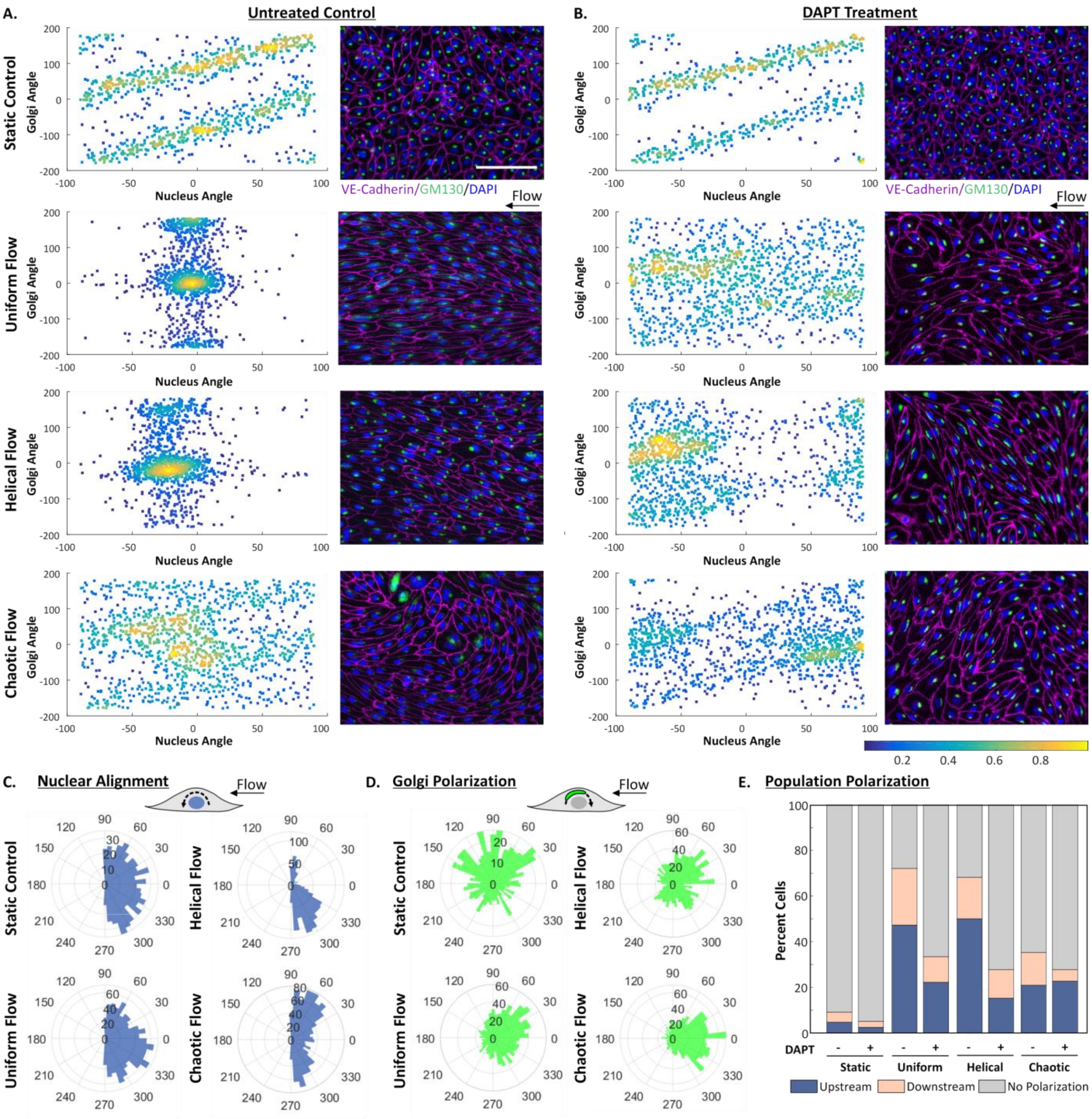


**Figure S9. Coupling of nuclear and golgi orientation and overall polarization at the single cell level under distinct SPF and following DAPT perfusion.** Density scatter plots of the nuclear and golgi orientation for single cells under distinct SPF, without (**A.**) or with (**B.**) DAPT treatment. Subcellular localization of hEC golgi complex (GM130, green stain) with respect to the cell (VE-Cadherin, magenta stain) and nucleus (DAPI, blue stain) are shown as well. All data was accumulated at 72h after perfusion at 20 dynes/cm$^2$. Control cells were preconditioned with flow for 48h, followed by vehicle treatment and the NOTCH1-inhibited cells were similarly preconditioned and treated with DAPT at 10 µM for 24h under flow. **C.** Nuclear alignment and (**D.**) golgi polarization distributions are shown under the same DAPT-treatment conditions. **E.** Population polarization was determined based on single-cell polarity classification, with or without DAPT treatment (for all panels: n =3, 1250 cells/condition). Scale: 0.1 mm.

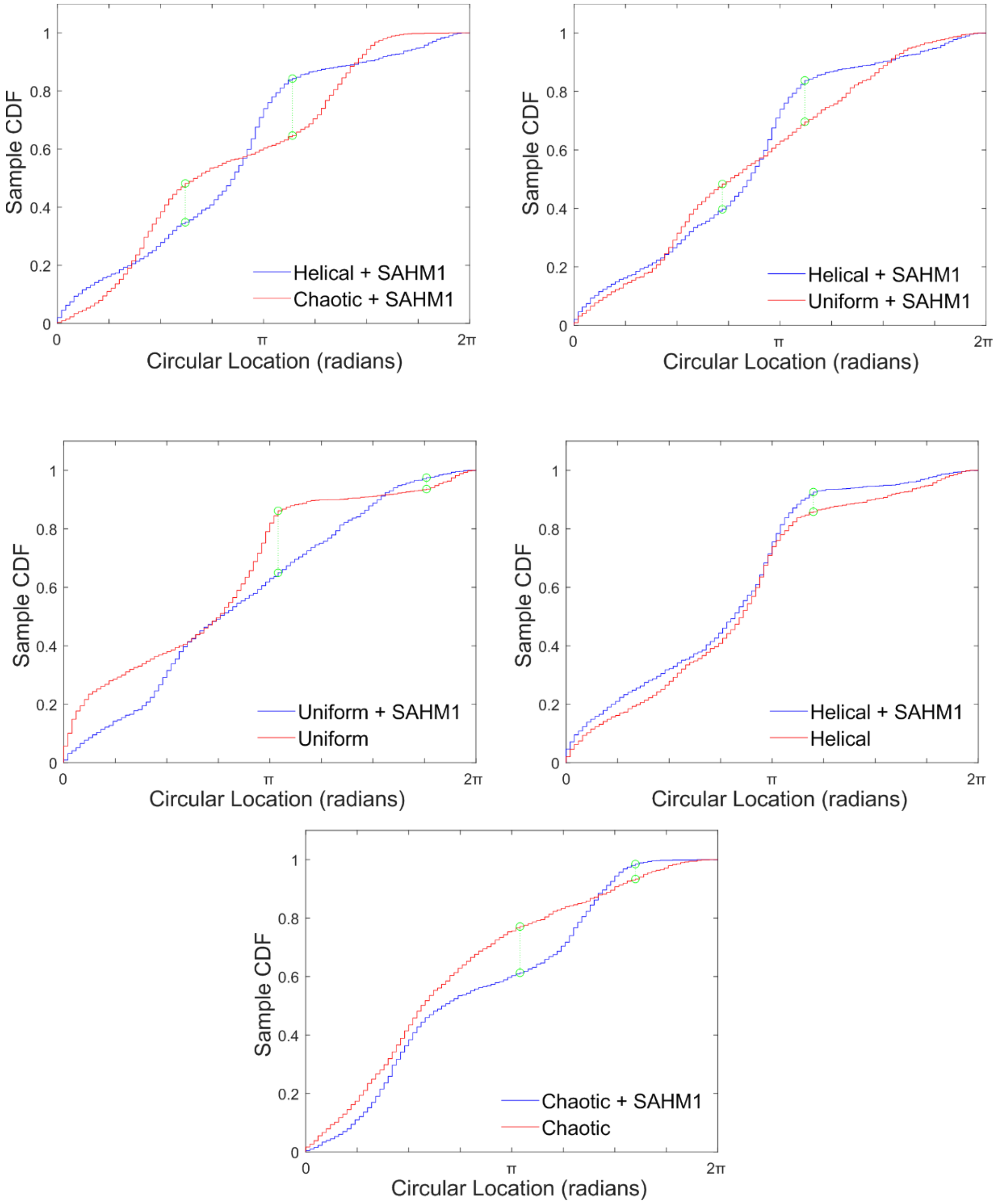


**Figure S10. Cumulative distribution functions and Kuiper tests of relative golgi angles following SAHM1 perfusions.** Relative golgi angle to the nucleus was calculated for each cell under each of the conditions shown above and all relative angles were used to generate cumulative distribution functions (CDF) over the spectrum of circular locations. These CDF were then used in the Kuiper statistical test to assess statistical difference in the angular distributions each of the condition pairs shown above. Test statistics $D^+ + D^-$ (absolute sizes of the most positive and most negative differences between the CDFs) are shown in green, and p-values are summarized in Supplemental Table III. Data pooled from ~1250 cells/conditions, n =2 experiments)

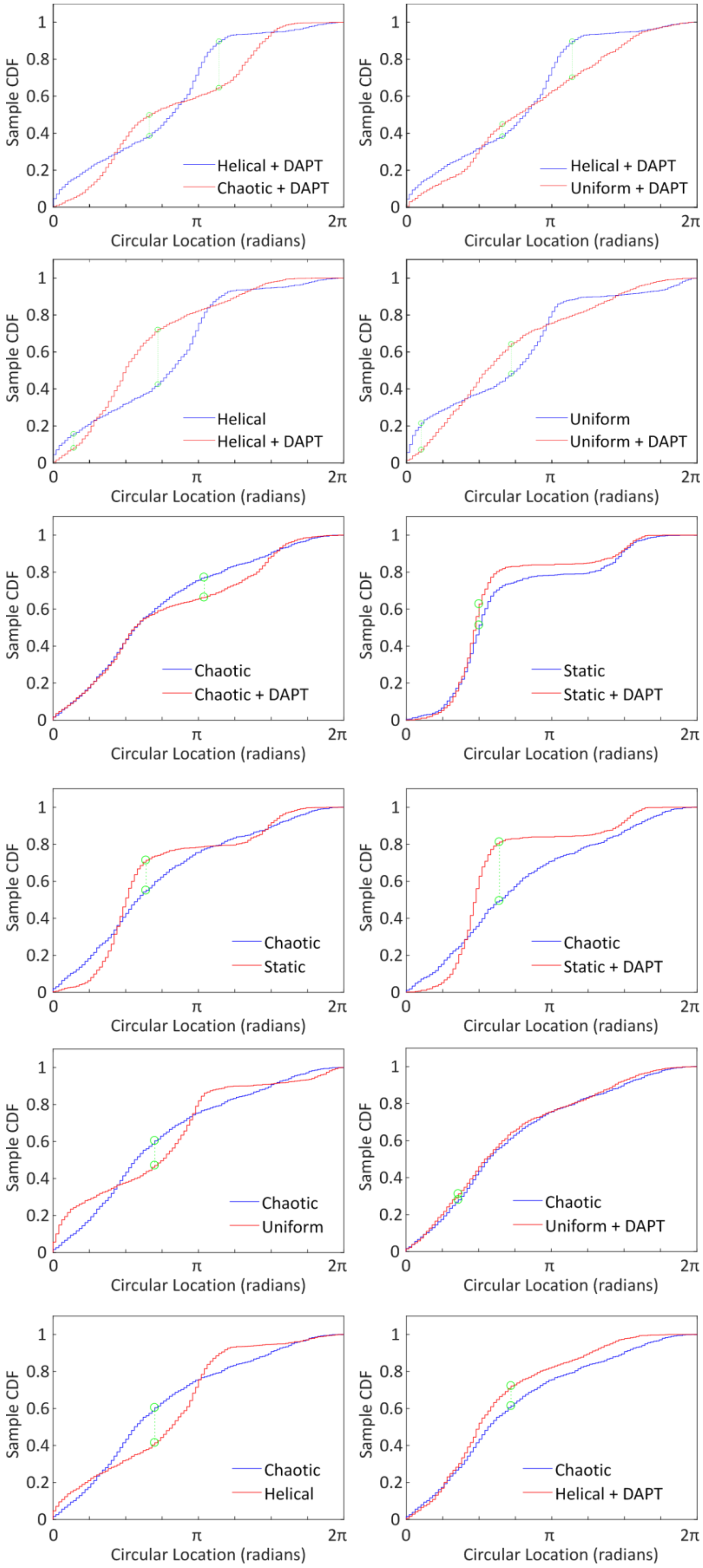

Sample CDF
Circular Location (radians)
0
π
2π
0
0.2
0.4
0.6
0.8
1
Helical + DAPT
Chaotic + DAPT
Helical + DAPT
Uniform + DAPT
Helical
Helical + DAPT
Uniform
Uniform + DAPT
Chaotic
Chaotic + DAPT
Static
Static + DAPT
Chaotic
Static
Chaotic
Static + DAPT
Chaotic
Uniform
Chaotic
Uniform + DAPT
Chaotic
Helical
Chaotic
Helical + DAPT

**Figure S11. Cumulative distribution functions and Kuiper tests of relative golgi angles following DAPT treatment.** Relative golgi angle to the nucleus was calculated for each cell under each of the conditions shown above and all relative angles were used to generate cumulative distribution functions (CDF) over the spectrum of circular locations. These CDF were then used in the Kuiper statistical test to assess statistical difference in the angular distributions each of the condition pairs shown above. Test statistics $D^+ + D^-$ (absolute sizes of the most positive and most negative differences between the CDFs) are shown in green, and p-values are summarized in Supplemental Table III. Data pooled from ~1250 cells/conditions, n =3 experiments)

### Supplementary Tables

##### Supplemental Table I. Statistical measurements and tests for cellular orientation

| | Static | Uniform | Helical | Chaotic |
|---|---|---|---|---|
| Mean | -4.266 | -3.462 | -18.799 | -2.835 |
| Median | -4.548 | -3.535 | -19.619 | -4.418 |
| Variance | 20.156 | 1.223 | 1.572 | 20.382 |
| Standard Dev. | 48.059 | 11.840 | 13.422 | 48.328 |
| Kruskal–Wallis H test of the distribution median angle against static (p- value) | N/A | 0.6726 | <0.001 | ns |
| Kuiper Test of population distribution against static (p-value) | N/A | <0.001 | <0.001 | ns |

##### Supplemental Table II. Statistical measurements and tests golgi orientation

| | Static | Uniform | Helical | Chaotic |
|---|---|---|---|---|
| Mean | 86.840 | 18.358 | -7.163 | 19.393 |
| Median | 74.745 | 10.305 | -10.008 | 287.108 |
| Variance | 51.944 | 42.236 | 36.299 | 48.358 |
| Standard Dev. | 77.151 | 69.569 | 64.495 | 74.441 |
| Kruskal–Wallis H test of the distribution median angle against static (p- value) | N/A | <0.001 | <0.001 | <0.001 |
| Kuiper Test of population distribution against static (p-value) | N/A | <0.001 | <0.001 | <0.001 |

##### Supplemental Table III. Statistical analyses of relative golgi angles under distinct SPF conditions

| | Kruskal–Wallis H test of the distribution median (p-value) | Kuiper Test of population distribution (p-value) |
|---|---|---|
| **DMSO Control** | | |
| Static vs. Chaotic | 0.0206 (*) | <0.001 (***) |
| Uniform vs. Chaotic | 0.0027 (**) | <0.001 (***) |
| Helical vs. Chaotic | <0.001 (***) | <0.001 (***) |
| **SAHM1 Treatment** | | |
| Uniform (+SAHM1) vs. Uniform (-SAHM1) | 0.0903 (ns) | <0.001 (***) |
| Helical (+SAHM1) vs. Helical (-SAHM1) | 0.3371 (ns) | 0.1 (ns) |
| Chaotic (+SAHM1) vs. Chaotic (-SAHM1) | <0.001 (***) | <0.001 (***) |
| Helical (+ SAHM1) vs. Uniform (+SAHM1) | 0.0083 (**) | <0.001 (***) |
| Helical (+SAHM1) vs. Chaotic (+SAHM1) | <0.001 (***) | <0.001 (***) |
| Static (+ SAHM1) vs. Static (- SAHM1) | 0.4398 (ns) | <0.001 (***) |
| **DAPT Treatment** | | |
| Uniform (+DAPT) vs. Uniform (-DAPT) | 0.0018 (**) | <0.001 (***) |
| Helical (+DAPT) vs. Helical (-DAPT) | <0.001 (***) | <0.001 (***) |
| Chaotic (+ DAPT) vs. Chaotic (- DAPT) | 0.0375 (*) | <0.001 (***) |
| Helical (+ DAPT) vs. Uniform (+DAPT) | 0.0164 (*) | <0.001 (***) |
| Helical (+DAPT) vs. Chaotic (+DAPT) | <0.001 (***) | <0.001 (***) |

| | | |
|---|---|---|
| Static (+ DAPT) vs. Static (- DAPT) | 0.0867 (ns) | <0.001 (***) |
| Static (+ DAPT) vs. Chaotic (- DAPT) | 0.0024 (**) | <0.001 (***) |
| Uniform (+ DAPT) vs. Chaotic (- DAPT) | 0.7829 (ns) | 1 (ns) |
| Helical (+ DAPT) vs. Chaotic (- DAPT) | 0.4715 (ns) | <0.001 (***) |

Note: Statistical tests for circular distributions were adapted and implemented without the assumption of underlying von Mises distributions.

**Supplemental Table IV. Primer Sequences**

| Gene | Forward Primer Sequence | Reverse Primer Sequence |
|---|---|---|
| GAPDH | GCCGCATCTTCTTTTGCGTC | TACGACCAAATCCGTTGACTCC |
| NRARP | TGCAGAAATGGGGAGCACTT | ATAAAAAGCGACGGAGGGCT |
| HES1 | AGCACAGAAAGTCATCAAAGCC | ATGCCGCGAGCTATCTTTCT |
| HES4 | ACGCCCTCAGAAAAGAGAGC | AGCGCGGCCGTCACC |
| NOTCH1 | ACGTGGTGGACCGCAGA | GCTGGCACGATTTCCCTGAC |
| DLL4 | TGACCACTTCGGCCACTATG | CCCGAAAGACAGATAGGCTGTT |
